# FGF21 deletion mildly exacerbates hepatic dysfunction in MASH diet and alcohol fed rats

**DOI:** 10.1101/2024.03.06.583704

**Authors:** Peter Aldiss, Malte Hasle Nielsen, Hayley Burm, Denise Oró, Henrik H. Hansen, Michael Feigh, Matthew P Gillum

## Abstract

**Objective:** Fibroblast growth factor 21 (FGF21) is a hepatokine that improves dyslipidemia, steatosis, inflammation, and fibrosis. FGF21 analogues are in clinical development as treatments for metabolic and alcohol-associated liver disease, creating a need for new models to help understand FGF21 physiology and drug mechanisms of action. The aim of this study was to create and initially characterize the first FGF21 knockout (KO) rat line to validate its utility as a translational animal model that recapitulates human MASH and ALD, and to provide a resource for examining FGF21-related phenotypes that are more appropriate for the rat.

**Methods:** We generated an FGF21 KO rat model using CRISPR/Cas9 to insert an artificial STOP codon in exon 1 and exposed 6-month-old WT and KO rats to either chow (n=8 per genotype) or the GAN (Gubra Amylin NASH) diet (n=16 per genotype) for 12 weeks. We further evaluated alcohol drinking behavior and biochemistry in FGF21 KO and WT rats. In the diet model, we further analysed liver and blood biochemistry in addition to histopathological scoring of NAFLD activity score (NAS), fibrosis stage and the liver transcriptome for in-depth characterization of the model.

**Results:** Lack of endogenous FGF21 increased plasma transaminases, liver weight, and total levels of liver TG in GAN-fed FGF21 KO rats. FGF21 deletion also increased ALT in alcohol-fed FGF21 KO rats. However, in the GAN diet model, FGF21 KO had no impact on body weight, fat mass, glycaemic traits, MASH histological endpoints including hepatic steatosis, NAS score, lobular inflammation, ballooning degeneration or fibrosis stage after 12 weeks. Similarly, there was no effect of the loss of endogenous FGF21 on the liver transcriptome in response to GAN diet feeding. Finally, we demonstrate that endogenous FGF21 does not regulate drinking behaviour in rats.

**Conclusion:** FGF21 deficiency accelerates hepatic dysfunction in diet and alcohol-induced liver disease models in rats, providing support from a new species that FGF21 might be a hepatoprotective factor.

## 1. Intro

Obesity and type 2 diabetes are associated with multiple co-morbidities including excess lipid accumulation in the liver and the progression to metabolic dysfunction-associated fatty liver disease (MAFLD) [1]. MAFLD is the most common chronic liver disease in the western world and is a major risk factor for hepatocellular carcinoma and cirrhosis [1]. During this progression, MAFLD advances to metabolic dysfunction-associated steatohepatitis (MASH) where the liver is characterised by lobular inflammation, steatosis and ‘ballooning’, or hepatocyte injury [1]. At present, there are no FDA-approved treatments for MASH and clinically relevant models are sparse and, as such, there is a critical need for development of these models to support drug-development pipelines in the search for effective treatments.

Fibroblast growth factor 21 (FGF21) is a liver secreted hormone, or hepatokine, which is also expressed in adipose tissues. Originally identified as a ‘starvation’ factor, it is now clear that FGF21 has numerous physiological roles including the regulation of glucose and lipid metabolism, energy balance and appetite which are mediated through its binding to FGF receptor 1 (FGFR1) and the co-receptor β-klotho (KLB).

Mechanistically, FGF21 is a critical regulator of hepatic metabolism and plays a key role in the development of MAFLD [2]. FGF21 knockout mice develop significant hepatic steatosis in response to ketogenic, and amino acid deprived diets through impaired fatty acid oxidation and increased lipogenesis [3-5]. More broadly, FGF21 knockout mice are hyperinsulinaemic and develop attenuated liver insulin sensitivity in response to an obesogenic diet [6, 7]. Conversely, FGF21 overexpression, or treatment with FGF21, improves insulin sensitivity, reduces hepatic steatosis, increases thermogenic capacity of white and brown adipose tissues, and improves β-cell function [2]. Further, FGF21 analogues consistently reverse, or reduce MASH in genetic, diet and chemically induced pre-clinical models demonstrating the therapeutic potential of FGF21 analogues in resolving liver pathologies [8].

In humans, circulating FGF21 is increased in MAFLD and MASH, increasing with the degree of fibrosis and is an independent predictor of hepatic steatosis [9]. Circulating FGF21 is inversely correlated with insulin sensitivity and correlates positively with visceral fat mass and circulating ALT [9, 10]. Further, long-acting FGF21 analogues which are currently in clinical trials consistently report reductions in circulating triglycerides and cholesterol, hepatic triglycerides and potentially improve liver fibrosis [11, 12]. Despite this, the mechanisms underlying the role of FGF21 in MASH remain unclear.

Further, FGF21 is implicated in the genetics of alcohol intake and alcohol use disorder. Pre-clinical models demonstrate that FGF21 treatment can reduce alcohol preference, and intake, in mice and non-human primates where it acts through an amygdalo-striatal circuit [13]. As such, advanced models are now required to further understand FGF21 biology and its role in complex behaviours such as addiction.

To this end, and taking advantage of recent advancements in CRISPR/Cas9 technology, we capitalised on the translational value of rat models of liver disease and addiction to developed a novel, FGF21 knockout rat and determine its suitability as a genetic, physiological model for human MASH and alcohol use disorder.

## 2. Methods

### 2.1 Ethics statement

All in vivo experiments were conducted at the University of Copenhagen, Denmark under approval from the Danish Animal Experiments Inspectorate, Danish Ministry of Food, Agriculture and Fisheries (Permit #). Forty-eight animals (n=24 WT, n=24 KO) were randomized to either chow (n=8 per group, 3.22⍰kcal/g, Altromin 1324, Brogaarden, Hoersholm, Denmark) or the Gubra-Amylin MASH (GAN) diet (n=16 per group, 4.49⍰kcal/g, 40⍰kcal-% fat (of these 46% saturated fatty acids by weight), 22% fructose, 10% sucrose, 2% cholesterol; D09100310, Research Diets) for a period of 12 weeks. Rats were pair-housed, supplied with enrichment in a temperature-controlled environment (21-23°C) and maintained on a 12-h light-dark cycle (6:00 am - 6:00 pm) with ad libitum access to food and drinking water, unless otherwise specified. Animals were terminated by cardiac puncture under isoflurane anesthesia.

For alcohol studies, WT and KO rats (n=15 per group) were exposed to 10% ethanol for 3 weeks across 3 different drinking paradigms. In week 1, animals were given continuous access to water or 10% ethanol (two-bottle preference). In week 2, animals were exposed to the first a binge drinking paradigm and given continuous access to water with 10% ethanol for 24h on Mon, Wed, Fri (intermittent access two-bottle choice) In week 3, animals were exposed to a final binge drinking paradigm where they have continuous access to water and 10% ethanol was available for 2h periods, beginning 2h after the onset of the dark cycle when drinking behaviour peaks (drinking in the dark). Animals were terminated by cardiac puncture under isoflurane anesthesia.

### 2.2 Model generation

The FGF21 knockout rat model was developed by genOway (Lyon, France) on a Wistar background. CRISPR/Cas9 was used to insert an artificial STOP codon in exon 1, leading to production of an unstable mRNA and the 8 amino acids critical for FGF21 activity were replaced. All in-vivo experiments were performed in male rats aged around 6 months old (chow, n11=1116, 8 per genotype; GAN, n11=1132, 16 per genotype).

### 2.3 Plasma and liver biochemistry

Animals were fasted for 4 hours after which blood was sampled from the sublingual vein, kept on ice, and centrifuged (5 min, 4°C, 6000 g) to generate EDTA-stabilized plasma. Insulin, alanine aminotransferase (ALT), aspartate aminotransferase (AST), triglycerides (TG), total cholesterol (TC), and liver TG and TC were determined as described previously [14]. Plasma glucose was measured by glucometer (Roche, Basel, Switzerland) in animals following a brief 4h fast. FGF21 was measured in duplicate in EDTA plasma samples using the Mouse/Rat FGF-21 Quantikine ELISA Kit (Bio-Techne, MF2100) in accordance with the manufacturer’s instructions. For alcohol preference studies ALT was measured using the Rat ALT ELISA Kit (Abcam, Cambridge, UK) as per manufacturer’s instructions.

### 2.4 Liver histology

A liver biopsy section (≈200 mg) was dissected from the left lateral lobe at termination, fixed overnight in 4% paraformaldehyde, paraffin-embedded and sectioned (3 μm thickness). Liver sections were stained with hematoxylin-eosin (HE), picro-Sirius Red (PSR; Sigma-Aldrich, Brøndby, Denmark), anti-galectin-3 (cat. 125402, Biolegend, San Diego, CA), alpha-smooth muscle action (α-SMA, cat. Ab124964; Abcam, Cambridge, UK) or anti-type I collagen (Col1a1, cat. 1310-01, Southern Biotech, Birmingham, AL) according to standard procedures [14] [15].

### 2.5 Automated deep learning-based image analysis

Automated deep learning-based image analysis GHOST (Gubra Histopathological Objective Scoring Technique)[25] was used for objective histopathological scoring of steatosis, lobular inflammation, hepatocyte ballooning, and fibrosis staging according to the MASH Clinical Research Network (CRN) scoring system [16]. GHOST was further used to obtain histopathological scoring-associated variables, including fraction of lipid-laden hepatocytes (%), number of inflammatory foci (foci/ mm2), and percent proportionate area fibrosis. Quantitative histology was performed using Visiomorph digital imaging software (Visiopharm, Hørsholm, Denmark) for the determination of liver fat (HE-staining), fibrosis (PSR, Col1a1), inflammation (galectin-3), and alpha-smooth muscle actin (α-SMA) expressed relative (%) to total sectional area. Percent area of positive staining was multiplied with the corresponding total liver weight to give an estimate of total liver marker content (mg).

### 2.6 Liver RNASeq

Total RNA was isolated using RNeasy Minikit (Qiagen) according to the manufacturer’s protocol. Total RNA sequencing was performed by the Single-Cell Omics platform at the Novo Nordisk Foundation Center for Basic Metabolic Research. Libraries were prepared using Universal Plus Total RNAseq with NuQuant kit (Tecan) as recommended by the manufacturer. In brief, 250ng total RNA was fragmented followed by cDNA synthesis. Adaptors were ligated and after stand selection, the cDNA was depleted from rRNA using AnyDeplete probes (Tecan). Following the final bead clean-up, libraries were quantified with NuQuant, quality checked using a TapeStation instrument (Agilent Technologies) and subjected to 52-bp paired-end sequencing on a NovaSeq 6000 (Illumina). Differential gene expression analysis was conducted using the DESeq2 R-package. Gene set analysis was performed using the PIANO version 1.18.1 R-package and the Stouffer method. The Reactome pathway database [17] was used for gene annotation for gene set analysis. The Benjamini-Hochberg method (5% False Discovery Rate, FDR < 0.05) was used for multiple testing correction of P values.

### 2.7 Statistics

Except for RNA sequencing and deep learning-based image analysis, data were analyzed using GraphPad Prism v9.0.2 software (GraphPad, La Jolla, CA). Data are presented as mean ± standard error of mean (SEM). Ordinary one-way or two-way ANOVAs were used for all statistical comparisons (specified in figure legend), except for histological scoring (Fig. 2A-2E) where Mann–Whitney U test with Bonferroni correction was used. A P-value of <0.05 was considered statistically significant.

## 3. Results

### 3.1 FGF21 deficiency increases liver weight and plasma transaminases but not body weight or fat mass

FGF21KO rats were deficient in circulating FGF21 (Fig. 2A). Histological and transcriptional markers of MASH were evaluated in WT and FGF21 KO rats after 12 weeks of GAN diet and compared to WT and FGF21 KO rats fed chow diet for 12 weeks (see study outline in Fig. 1). At baseline, prior to GAN diet induction, levels of plasma biochemical markers (insulin, triglycerides, total cholesterol, ALT and AST) were similar between all groups (Suppl. Fig. 2A-2E). GAN diet feeding for 12 weeks or FGF21 deficiency did not lead to changes in terminal body weight (Fig. 2D). GAN diet increased body weight gain (Fig. 2E). GAN diet increased and hepatomegaly which was further exacerbated by FGF21 deficiency (Fig. 2F). Furthermore, iWAT, eWAT, and BAT mass was increased in response to GAN diet but not affected by FGF21 deficiency (Fig. 2G-2I). 4h fasted blood glucose levels were similar between all groups at week 11 (Fig. 2J), while 4h fasted plasma insulin levels were only lower in WT GAN group compared to WT chow (Fig. 2K). Terminal biochemistry analysis displayed a GAN-driven increase in total plasma cholesterol, ALT, AST and a decrease in plasma triglycerides (Fig. 2L-2O). FGF21 deficiency in the context of GAN diet feeding further increased plasma AST and trended towards increasing plasma ALT (p=0.05). GAN-diet feeding increased levels of liver triglycerides and total cholesterol in both WT and FGF21 KO rats.

**Figure 1.**
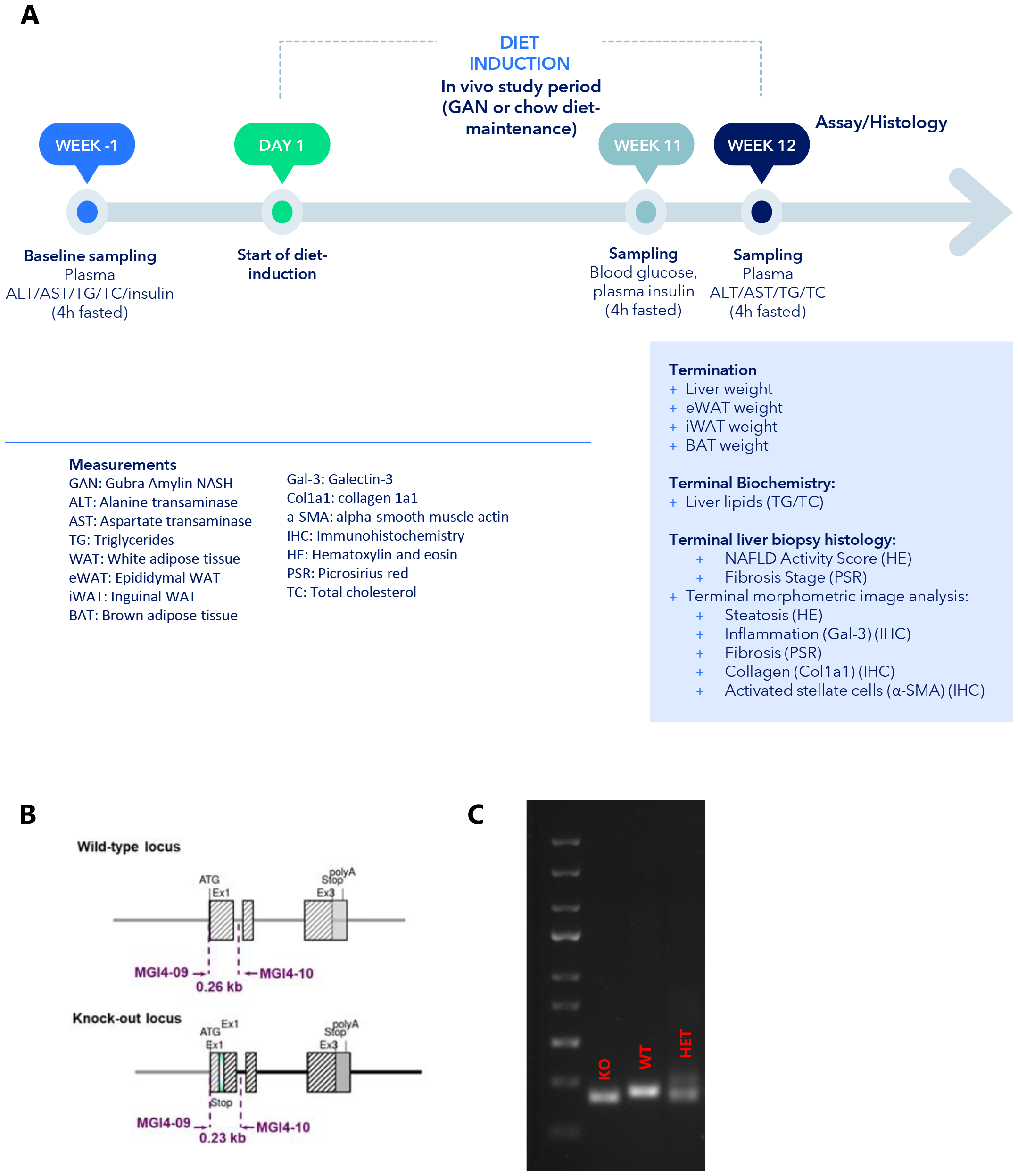
Overview of study design and endpoints **(A)** Overview of model generation by genOway (Lyon, France) using CRISPR/Cas9 to insert an artificial STOP codon in exon 1 **(B)** and example of genotyping **(C)**.

**Figure 2.**
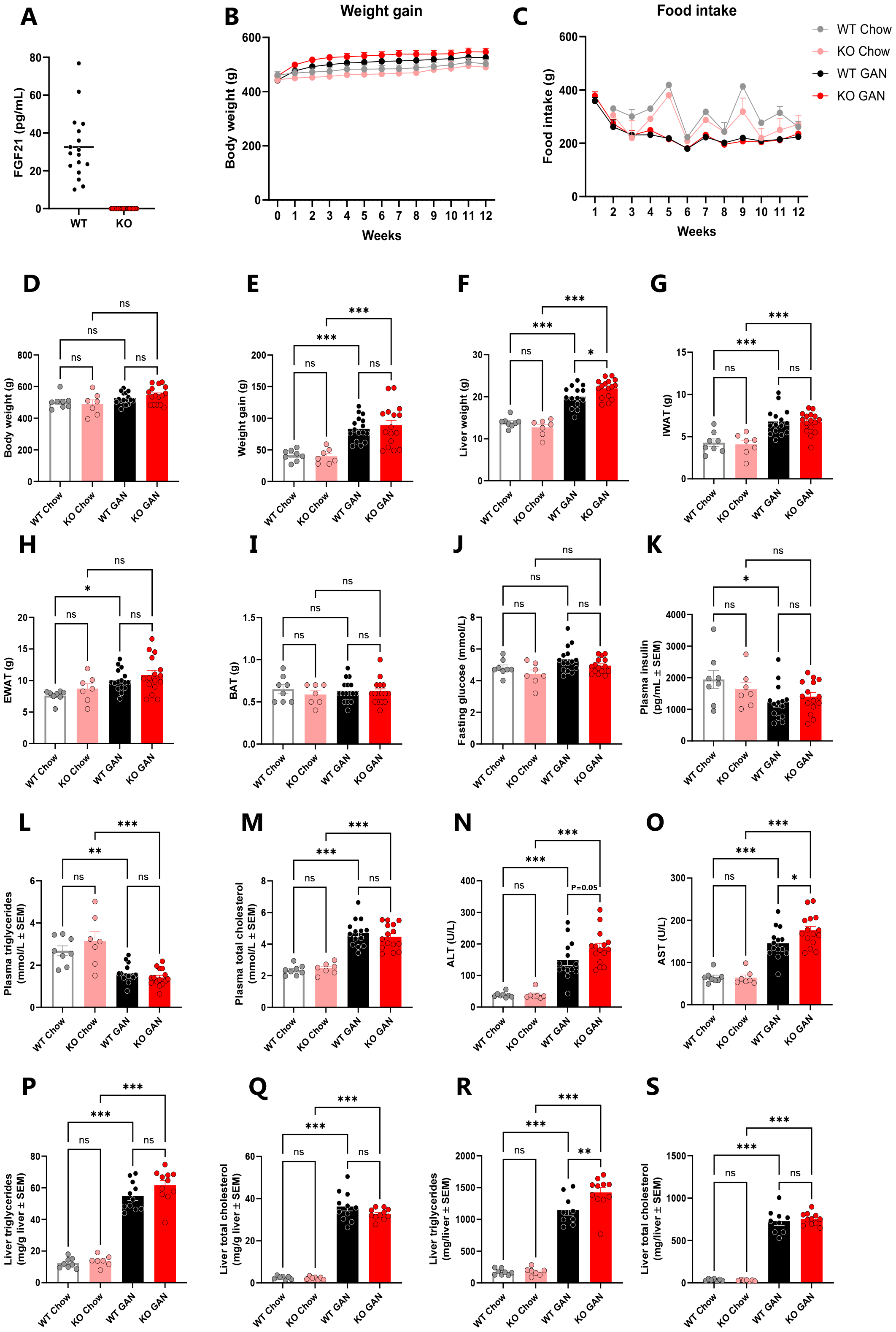
Effect of 12 weeks of GAN diet feeding on bogy weight, food intake, fat mass, metabolic and biochemical parameters in FGF21 KO rats. **(A)** Validation of circulating FGF21. **(B)** Body weight trajectories. **(C)** Food intake. **(D)** Terminal body weight. **(E)** 12-week body weight gain. **(F)** Terminal liver weight. **(G)** IWAT weight. **(H)** EWAT weight. **(I)** BAT weight. **(J)** 4h fasted blood glucose. **(K)** 4h fasted plasma insulin. **(L)** 4h fasted plasma triglycerides. **(M)** 4h fasted plasma total cholesterol. **(N)** 4h fasted plasma ALT. **(O)** 4h fasted plasma AST. **(P)** Liver triglyceriders. **(Q)** Liver total cholesterol. *n* =7–16 rats/group, *P < 0.05, **P < 0.01, ***P < 0.001 (A-E, Mann–Whitney U test with Bonferroni correction; F-J, Ordinary One-way ANOVA). GAN, Gubra Amylin MASH; WAT, white adipose tissue; IWAT, inguinal WAT; EWAT, epididymal WAT; BAT, brown adipose tissue; ALT, alanine transaminase; AST, Aspartate transaminase.

### 3.2 FGF21 deficiency does not worsen GAN driven hallmarks of MASH

Rats fed the GAN diet for 12 weeks exhibited moderate MAFLD (NAS of 2-5), with more than half of the animals featuring with NAS of ≥4 (Fig. 3A). Changes in NAS were primarily driven by increases in steatosis and lobular inflammation sore (Fig. 3C, 3D). There was no difference between WT and FGF21 KO rats. In both groups, only a few percentages of animals presented ballooning degeneration in response to GAN feeding (Fig. 3E). In support of changes in histopathological scores, quantitative markers of steatosis (% area of steatosis, Fig. 3F) and inflammation (% area of galectin-3, Fig. 3G) were increased in response to GAN feeding, but not affected by genotype status. 12 weeks of GAN diet feeding did not induce liver fibrosis as evident from fibrosis staging and corroborated by quantitative histological markers of fibrosis (PSR % area, Fig. 3H; Col1a1 % area, Fig. 3I). However, α-SMA, marker of hepatic stellate cell activation, was increased in the KO GAN group compared to KO Chow (Fig. 3J).

**Figure 3.**
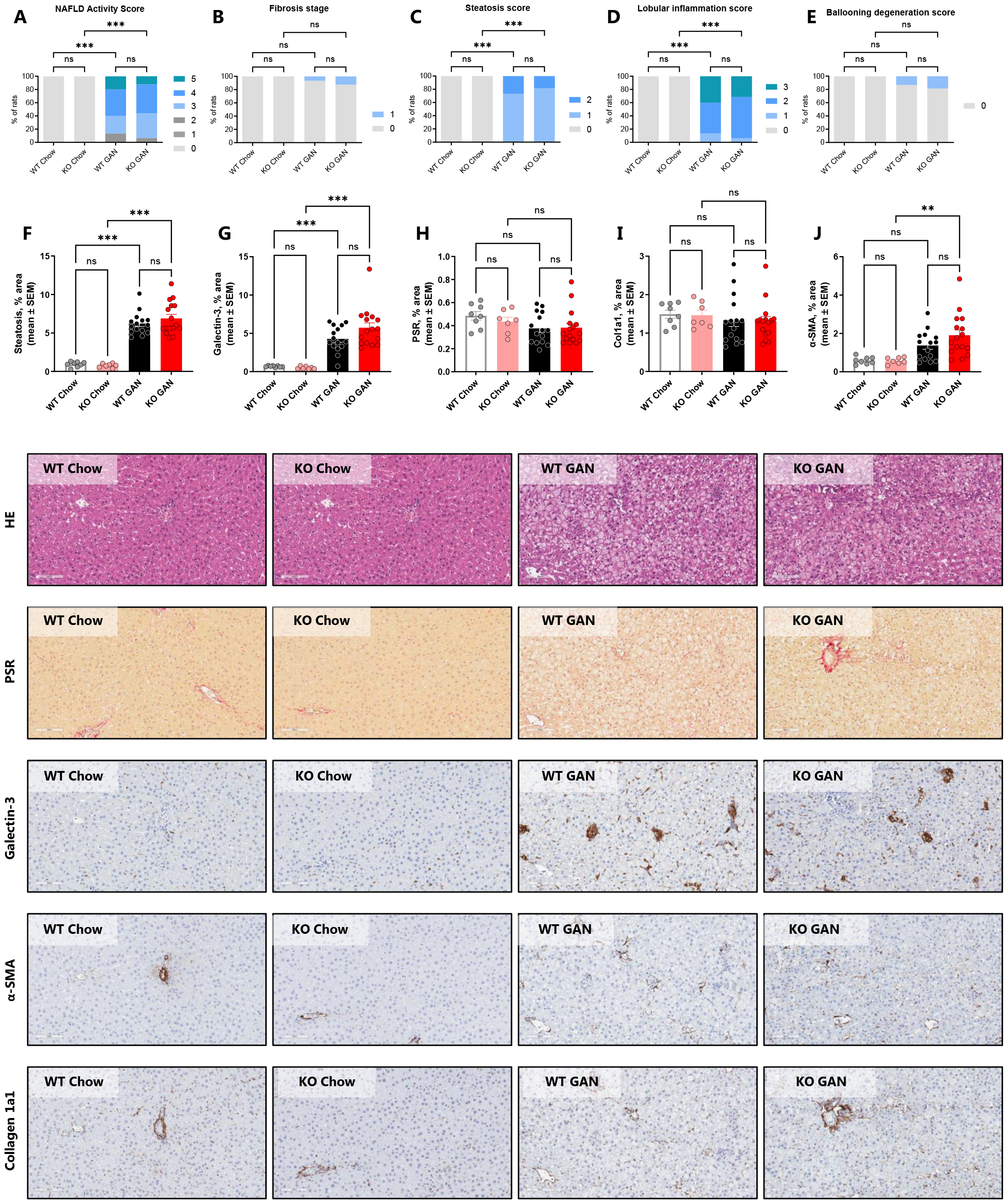
Effect of 12 weeks of GAN diet feeding and FGF21 deficiency on MAFLD Activity Score, fibrosis stage and quantitative histological markers. Histopathological scores were determined by Gubra Histopathological Objective Scoring Technique (GHOST) deep learning-based image analysis. **(A)** NAS. **(B)** Fibrosis stage. **(C)** Steatosis score. **(D)** Lobular inflammation score. **(E)** Ballooning degeneration score. Percent area of **(F)** steatosis, **(G)** Galectin-3, **(H)** PSR, **(I)** Collagen 1a1, and **(J)** α-SMA. *n* =7–16 rats/group, *P < 0.05, **P < 0.01, ***P < 0.001 (Ordinary One-way ANOVA). MAFLD, metabolic dysfunction-associated fatty liver disease, Gubra Amylin MASH; NAS, NAFLD Activity Score; GHOST, Gubra Histopathological Objective Scoring Technique; PSR, picrosirius red; α-SMA, alpha-smooth muscle actin.

### 3.3 Minimal changes in liver transcriptome signatures

A principal component analysis (PCA) of the 500 most variable genes displays clustering based on diet type along the PC1 axis, accounting for 86% of the variance (Fig. 4A). Genotype status had limited effect on the number of differentially expressed genes (DEGs) (Fig. 4B). The FGF21 KO chow group showed 21 regulated genes compared to WT chow, while there were no DEGs between FGF21 KO GAN and WT GAN. Although no MASH-associated candidate genes were significantly regulated, FGF21 status conferred minor trends in global gene expression (Fig. 4C).

**Figure 4.**
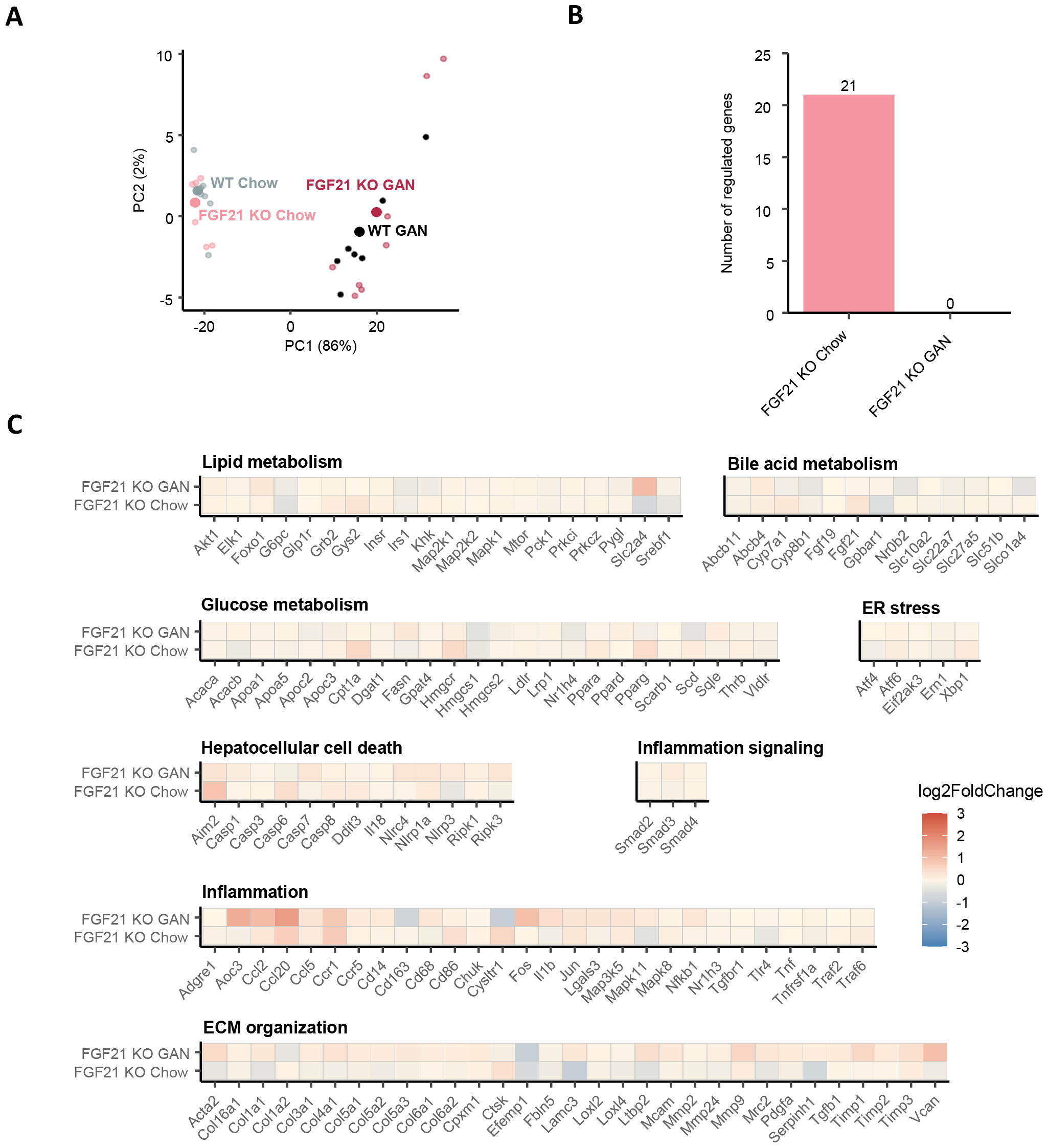
Transcriptional alterations in response to GAN diet feeding and FGF21 deficiency. **(A)** Principal component analysis (PCA) of samples based on top 500 most variable gene expression levels. **(B)** Total number of genes differentially expressed versus WT Chow and WT GAN, respectively. **(C)** Heatmaps displaying regulations in MASH and fibrosis-associated candidate genes (log2-fold change compared with respective WT group). Blue color gradients indicate downregulation of gene expression, red color indicate upregulation of gene expression. *n* =7–8 rats/group. GAN, Gubra Amylin MASH

### 3.4 Endogenous FGF21 does not regulate alcohol preference or binge drinking in rats

Finally, we explored the role of FGF21 to regulate behavior in the rat by studying its role in alcohol preference and in binge-drinking paradigms in separate cohorts. However, rats lacking endogenous FGF21 did not exhibit increased preference for alcohol during continuous access to either water or 10% ethanol (Supp Fig. 3A) and did not show increased alcohol consumption during two separate binge-drinking paradigms (Supp Fig. 3B-E). Despite this, FGF21 KO rats exhibited a significant increase in ALT during this period (Supp Fig. 3F).

## 4. Discussion

MASH is an advanced stage liver disease with no FDA-approved treatments. It is evident that the liver hormone FGF21 plays a role in both the development, and treatment of MASH with FGF21 analogues demonstrating positive results in clinical trials. However, the mechanisms underlying the role of FGF21 in liver disease remain unclear.

Here, we capitalised on the advancements of the gene-editing technology Crispr/Cas9, and the potential utility of rat models, by generating a novel FGF21 knockout rat model which could recapitulate the hallmarks of human disease when combined with the GAN diet. We demonstrate that FGF21 deficiency in the context of GAN diet feeding exacerbated hepatomegaly and plasma transaminases. However, both WT and FGF21 knockout rats display comparable weight curves and excess fat mass on a GAN diet with no difference in FGF21 knockout animals. GAN diet feeding increased MAFLD-associated plasma and liver markers (plasma TC/ALT/AST, liver TG/TC), with an additional worsening on plasma ALT/AST exerted by FGF21 deficiency. GAN diet consistently drove a liver phenotype characterised by increased liver steatosis and inflammation, evident from histopathological scores (NAS, steatosis score, lobular inflammation score) and supported by quantitative image analysis (steatosis % area, Galectin-3). There was no difference between WT and KO for any of the histological endpoints. In concert, transcriptional gene signatures clearly clustered according to diet with no additional effect of genotype status. However, GAN diet was not sufficient to drive advanced liver pathology with fibrosis.

Whilst mice are commonly used models to study MAFLD and MASH, primarily due to the genetic toolkits available, fibrosis models were originally developed in rats as they develop more robust fibrosis and are more susceptible to the histological hallmarks of MASH in response to diet [18, 19]. FGF21 knockout mice on a high-fat diet exhibit a worsening of metabolic liver injury, hepatic steatosis and MASH-HCC transition [20, 21]. Here, we describe a mildly exacerbated liver phenotype compared to wild-type animals, however, without displaying a worsening on histopathological endpoints. Although FGF21 analogues consistently reverse or reduce MASH in mouse models and FGF21 reduces body weight [22] and lower blood glucose in rats [8, 23], endogenous FGF21 in the current study design is not sufficient to drive the full-blown MASH with fibrosis.

The reasons for this are not clear, though it could be speculated that extending the diet-induction period could give rise to liver fibrosis. Indeed, mice develop fibrosing MASH after ≥28 weeks of GAN diet feeding [24, 25]. As such, it is possible that 12 weeks was insufficient time for rats to develop an advanced liver phenotype. Although, as discussed above, FGF21 KO mice exhibit exacerbated liver phenotypes in response to multiple different diets and it would have been expected, based on current evidence, that an FGF21 deficient rat would develop exacerbated liver phenotypes even over 12 weeks of exposure to the GAN diet. Similarly, it is possible that utilising younger animals may have yielded a more pronounced phenotype. Typically, mouse and rat models of diet-induced obesity commence between 6-10 weeks of age, around adolescence, as this is considered optimal in terms of weight gain, and often produces an experimentally clear metabolic phenotype in as little as 8-10 weeks. Given our aim was to develop a translationally relevant model that mimicked the hallmarks of human MASH, we commenced a MASH-inducing GAN diet in 24-week-old adult rats which, comparatively, will exhibit a lesser capacity for weight gain and therefore metabolic deterioration. However, it could be argued that, if endogenous FGF21 was an important regulator of liver pathology in rats, the age that animals commenced a diet should be less decisive. Indeed, FGF21 KO mice exhibit a subtle, but significant increase in body weight by 24 weeks on standard chow [3] which was not evident here and which one would expect to be exacerbated during high-fat feeding.

Finally, we show that endogenous FGF21 does not regulate drinking behaviour in rats. This in despite of a wealth of evidence implicating the beta-klotho-FGF21 pathway in the genetics of alcohol-use disorder [26, 27], demonstrating that FGF21 KO mice (global, liver and CNS-specific) drink more ethanol in both two-bottle preference, and binge drinking paradigms whilst FGF21 treatment reduces drinking in mice and non-human primates [13, 26]. Again, why endogenous FGF21 does not regulate drinking in the rat is unclear. There are strain-specific differences in alcohol intake, and susceptibility to ‘addiction’ but the Wistar rat, which we used as background, is commonly used due to their high alcohol preference and will choose alcohol over social interaction [28]. Importantly, the Wistar strain is also the most susceptible to MASH [29] and thus represents the best background strain to investigate the role of FGF21 in rats.

## 5. Conclusion

Through in-depth molecular, histological and behavioural characterisation we demonstrate that endogenous FGF21 mildly exacerbates hepatomegaly and biochemical markers of hepatic dysfunction in response to GAN diet and alcohol in rats.

## CRediT authorship contribution statement

**Conceptualization:** PA, MHN, DO, HHH, MF, MPG**; Data curation:** PA, MHN**; Formal analysis:** PA, MHN**; Funding acquisition:** PA, MHN, DO, HHH, MF, MPG**; Investigation**: PA, MHN, HB**; Methodology:** PA, MHN, DO, HHH, MF, MPG**; Project administration:** PA, MHN, MPG**; Resources:** DO, HHH, MF, MPG**; Software:** MHN, DO, HHH, MF**; Supervision:** PA, MHN, DO, MPG**; Visualization:** PA, MHN**; Writing - original draft:** PA, MHN**; Writing - review & editing:** PA, MHN, HB, DO, HHH, MF, MPG

## Financial support

PA was supported by the Danish Diabetes Academy (PD005-20), MHN was supported by Innovation Fund Denmark (0153-00178B). The Novo Nordisk Foundation Center for Basic Metabolic Research is an independent Research Center, based at the University of Copenhagen, Denmark, and partially funded by an unconditional donation from the Novo Nordisk Foundation (www.cbmr.ku.dk) (grant number NNF18CC0034900).

## Declaration of competing interest statement

M.H. Nielsen, D. Oró, H.H. Hansen, and M. Feigh are employed by Gubra. M.P. Gillum is employed by Novo Nordisk. None of the other authors has any conflicts of interest, financial or otherwise, to disclose.

## Acknowledgements

We acknowledge The Single-Cell Omics platform at the Novo Nordisk Foundation Center for Basic Metabolic Research (CBMR) and Maja Worm Andersen and Helene Ægidius at Gubra for the technical expertise and support with bioinformatic analyses.

---

**Supplementary figure 1.**
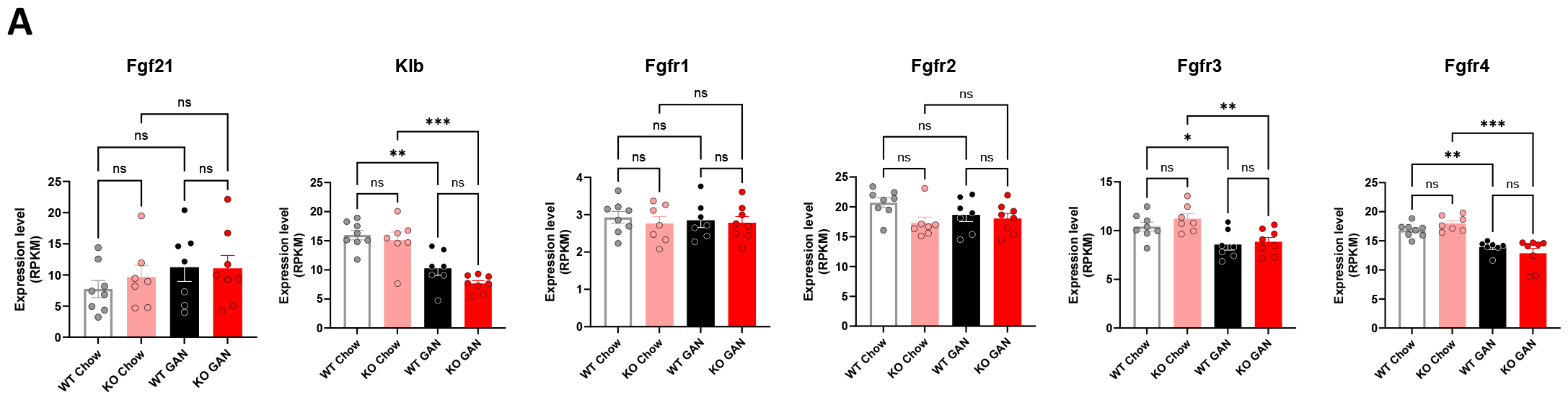
Hepatic mRNA expression of fibroblast growth factor 21 (Fgf21), b-klotho (Klb), FGF receptor 1 (Fgfr1), FGF receptor 2 (Fgfr2), FGF receptor 3 (Fgfr3), and FGF receptor 4 (Fgfr4). **P < 0.01, ***P < 0.001 (One-way ANOVA). n =7–8 rats/group.

**Supplementary figure 2.**
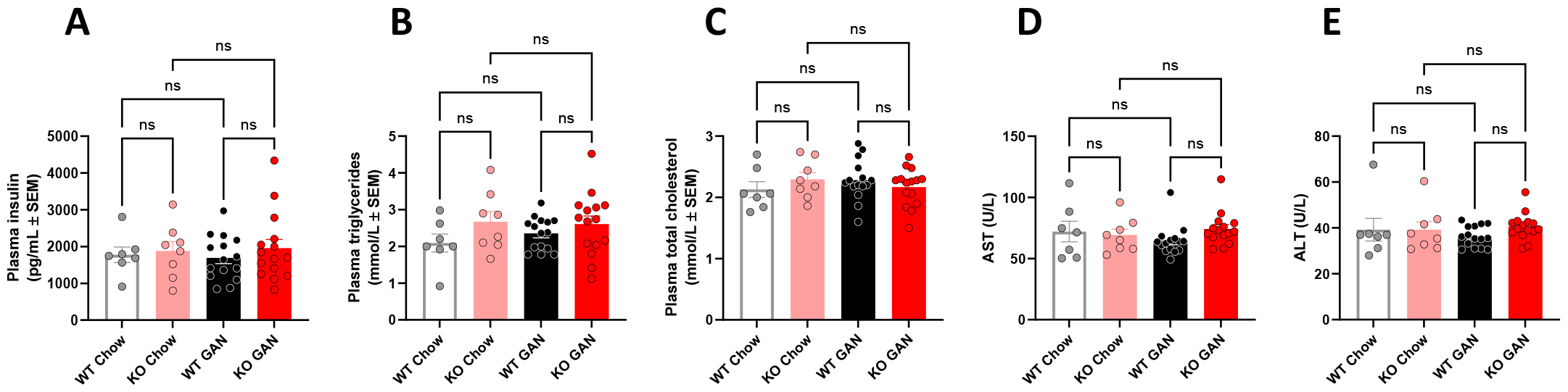
Baseline levels, prior to GAN diet induction, of plasma biochemical markers, 4h fasted. **(A)** Plasma insulin. **(B)** Plasma triglycerides. **(C)** Plasma total cholesterol. **(D)** Plasma ALT. **(E)** Plasma AST. *n* =7–16 rats/group, P > 0.05 (Ordinary One-way ANOVA).

**Supplementary figure 3.**
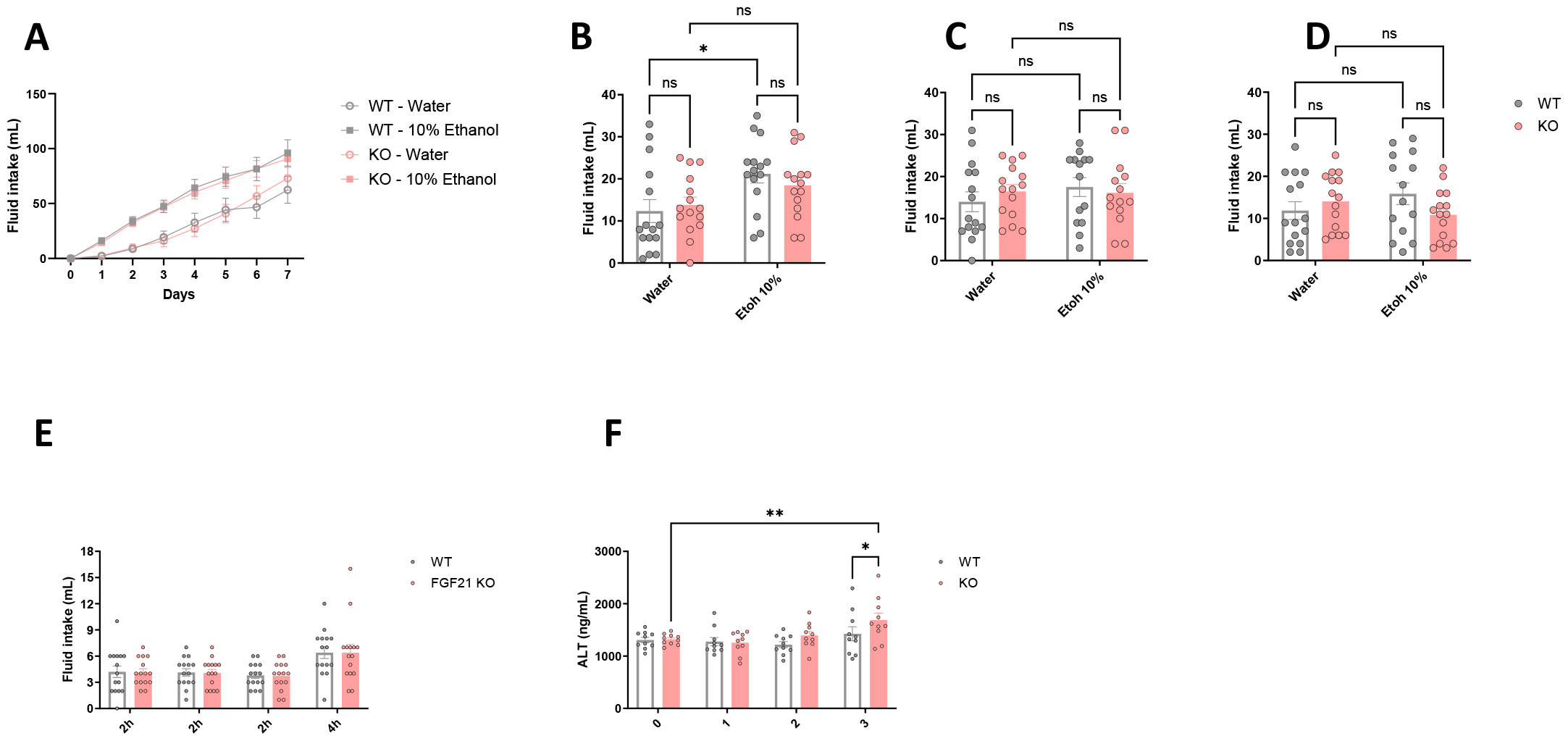
Drinking behaviour of FGF21 KO rats during (A) continous alcohol preference (B-D) Intermittent Access Two Bottle Choice days 1, 2 and 3 and (E) Drinking in the Dark studies. (F) Weekly changes in circulating ALT. *P < 0.05, **P < 0.01 (Two-way ANOVA). For (E) and (F), lines are only shown for significant comparisons. n =15 rats/group.

